# An Emergent Neural Coactivity Code for Dynamic Memory

**DOI:** 10.1101/776195

**Authors:** Mohamady El-Gaby, Hayley M Reeve, Vítor Lopes-dos-Santos, Natalia Campo-Urriza, Pavel V Perestenko, István Lukács, Ole Paulsen, David Dupret

## Abstract

Coincidental spike discharge amongst distributed groups of neurons is thought to provide an efficient mechanism for encoding percepts, actions and cognitive processes^1–3^. Short timescale coactivity can indeed bind neurons with similar tuning, giving rise to robust representations congruent with those of the participating neurons^4–6^. Alternatively, coactivity may also play a role in information processing through encoding variables not represented by individual neurons. While this type of emergent coactivity-based coding has been described for physically well-defined variables, including percepts and actions^7–10^, its role in encoding abstract cognitive variables remains unknown. Coactivity-based representation could provide a flexible code in dynamic environments, where animals must regularly learn short-lived behavioural contingencies. Here, we tested this possibility by training mice to discriminate two new behavioural contingencies every day, while monitoring and manipulating neural ensembles in the hippocampal CA1. We found that, while the spiking of neurons within their place fields is organised into congruent coactivity patterns representing discrete locations during unsupervised exploration of the learning enclosure, additional neurons synchronised their activity into spatially-untuned patterns that discriminated opposing learning contingencies. This contingency discrimination was an emergent property of millisecond timescale coactivity rather than the tuning of individual neurons, and predicted trial-by-trial memory performance. Moreover, optogenetic suppression of plastic inputs from the upstream left CA3 region during learning selectively impaired the computation of contingency-discriminating, but not space-representing CA1 coactivity patterns. This manipulation, but not silencing the more stable right CA3 inputs, impaired memory of the contingency discrimination. Thus, the computation of an emergent, coactivity-based discrimination code necessitates plastic synapses and supports dynamic, two-contingency memory.

---

To study the role of emergent coding in dynamic memory, we established a behavioural paradigm in which we trained mice to acquire two new stimulus-response-outcome associations (behavioural contingencies *X* and *Y*; Fig. 1a) within one newly encountered spatial configuration every day. At the start of each day, a learning enclosure was defined by a new spatial topology and position of two sets of LED wall-displays and two liquid dispensers (Fig. 1b and Supplementary Fig. 1). Mice initially explored the learning enclosure with each set of LEDs active in turn, as well as another (control) enclosure without any explicit task (Fig. 1c and Supplementary Fig. 1a). Next, we loaded both dispensers with either a rewarding (sucrose) or aversive (quinine) solution such that the identity of the drops to be simultaneously delivered at each port was non-matching, contingent upon the active LEDs, and signalled by the same auditory tone across both contingencies (Fig. 1a,b). Animals experienced alternating blocks of *X* and *Y* contingency trials, with one LED set active in a given block (Fig. 1c). Mice rapidly developed an efficient approach response to preferentially collect the drop at the correct (sucrose) dispenser over the incorrect (quinine) dispenser, and therefore demonstrated successful learning of the two LED-defined contingencies (Fig. 1d,e and Supplementary Fig. 1b). Memory of the learned contingencies was subsequently tested in a probe session, where the tone was presented without drop delivery, while pseudo-randomly switching between the two sets of LEDs (Fig. 1c) wherein mice continued to prefer the correct dispenser (Fig. 1f and Supplementary Fig. 1c). Thus, mice successfully learned to discriminate two new task contingencies each day, providing a paradigm to study the neural substrates of flexible memory in dynamic environments.

**Figure 1.**
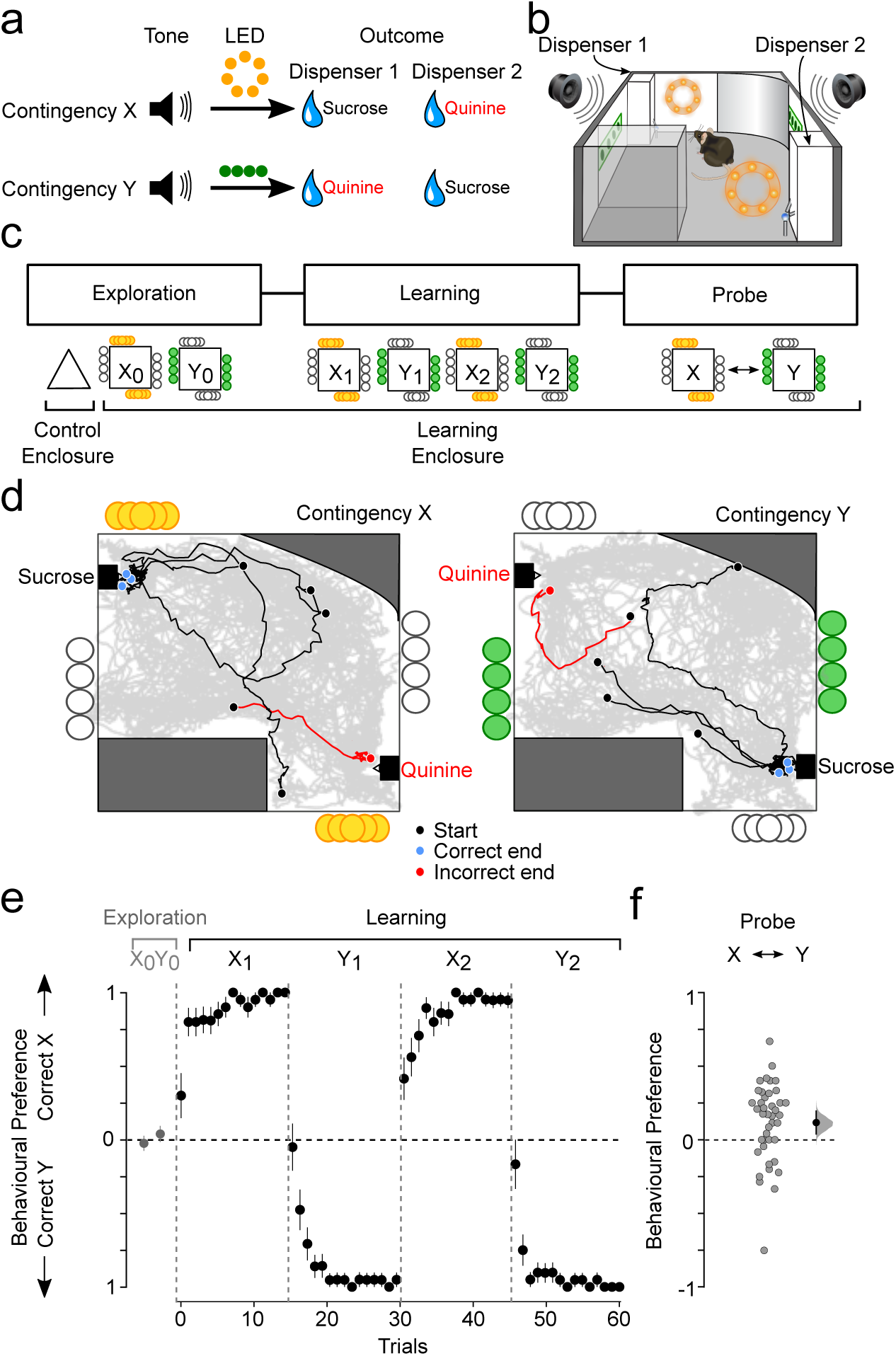
Mice perform flexibly on a one-day-two-contingency memory task. **a)** The two-learning-contingency layout. A tone signalled that both outcome dispensers deliver a liquid drop, the identity of which (sucrose versus quinine) depends on the active LED display. **b)** Schematic of an example learning enclosure. **c)** The three-stage task structure. Tone-defined trials occurred in Learning and Probe sessions, with the liquid drop outcomes only delivered during Learning. **d)** Example animal paths during trials in contingency *X* and contingency *Y* (correct path: black; incorrect path: red), overlaid on the overall animal path (grey) for one learning session. Black and blue/red circles represent path starting and correct/incorrect ending points, respectively. **e)** Behavioural preference for contingency-defined correct dispensers during Exploration and Learning (n=42 days from 13 mice). **f)** Behavioural performance during memory probe test, showing that animals preferred the correct dispenser during tone in each alternating contingency.

We monitored hippocampal CA1 neuronal ensembles throughout the exploration, learning and probe stages to investigate whether an emergent coactivity code develops in our two-contingency discrimination task. The short (25ms) timescale coactivity patterns^11^ detected in the learning enclosure (Fig, 2a) differed significantly from those extracted in the control enclosure (Fig. 2b), showing their spatial context-selective expression. In addition, we isolated learning enclosure coactivity patterns that were as highly selective for each contingency (i.e., *X* versus *Y*) as they were for the enclosure (Fig. 2b,c). To investigate the functional significance of such patterns, we compared them to a matched group of patterns with high between-contingency similarity (Fig. 2b,c). We refer to these learning-enclosure patterns of neuronal coactivity as contingency-discriminating and contingency-invariant patterns, respectively. Neurons that contributed the most to coactivity patterns are henceforth referred to as “member” neurons. Strikingly, member neurons of contingency-discriminating patterns were not individually contingency selective (Fig. 2d). Moreover, such coactivity-based discrimination could not be explained by differences in temporal firing properties of individual member neurons between contingencies (Supplementary Fig. 2a-c). Contingency-discrimination was instead seen at the level of temporal correlations between spike trains of such neurons (Supplementary Fig. 2d,e). Thus, we identify an emergent, neural coactivity-based discrimination of behavioural contingencies in the hippocampal CA1 during flexible learning.

**Figure 2.**
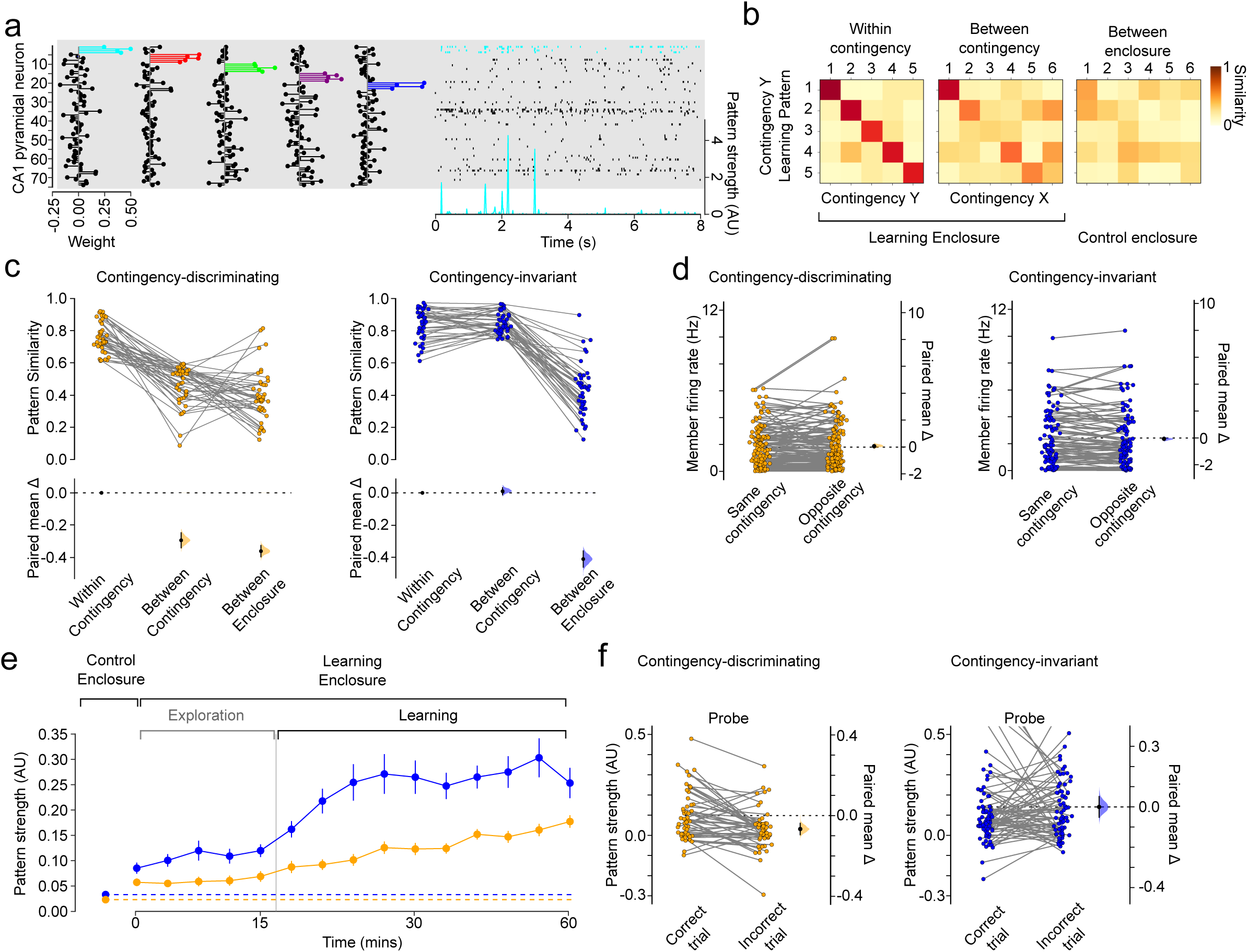
Emergent CA1 coactivity patterns discriminate task contingencies. **a)** Example CA1 coactivity patterns detected in the learning enclosure and represented as weight vectors (left) where the weight of each neuron in a given vector indicates the contribution of that neuron spiking to the coactivity defining that pattern^11^; neurons with a weight above 2 standard deviations of a given normalised vector were defined as member neurons (coloured) of that pattern. Projecting such vectors on neuron spike trains (right raster plot) monitored in any task session allowed tracking the timecourse of each pattern’s strength (e.g., light-blue peaks represent coactivity strength of the left most pattern, with the member neuron spiking shown in light-blue on the raster plot). **b)** Example similarity matrices of patterns detected in the learning enclosure with contingency *Y*, compared to patterns detected in subsequent sessions with the same (within contingency; left panel) or the other (between contingency; middle panel) contingency, or to patterns detected in the control enclosure (between enclosure; right panel). Overall pattern weight vector cosine similarities: within-contingency=0.72±0.01; between-contingency=0.66±0.01; between-enclosure=0.40±0.01; ANOVA with post-hoc Tukey-HSD test: P<0.001 for all comparisons; n=140 and 144 total patterns extracted in contingencies *X* and *Y* respectively; 5.0±0.3 total neurons per pattern. **c)** Across-condition similarity for contingency-discriminating and contingency-invariant patterns. Data Analysis with Bootstrap-coupled Estimation (DABEST^22^ plots are used from here onwards to plot data against a mean (or paired mean) difference between the two conditions (right y-axis) and to compare this difference against zero using the bootstrapping generated 95% confidence intervals (black error-bar). A kernel density estimate (shaded area) of the bootstrapping-generated resampled distribution is overlaid on the error-bar for clarity. **d)** Average firing rate of contingency-discriminating and contingency-invariant member neurons. **e)** Time-course of pattern strength. Dashed lines represent strength in Control enclosure. **f)** Pattern strength during probe trials, but before animal’s choice.

We next asked whether contingency-discriminating coactivity patterns are related to task performance. When we tracked the strength of each pattern over time (Fig. 2a) we found that contingency-invariant patterns began increasing in strength during the initial exploration of the new learning enclosure on each day, before animals experienced task contingencies; their strength further increased and subsequently plateaued during learning (Fig. 2e). Conversely, contingency-discriminating coactivity was stable during exploration but increased as animals learned task contingencies (Fig. 2e). Coactivity pattern strengthening during learning reflected increased coincidental spiking rather than member neuron firing rate changes (Supplementary Fig. 2f-i). Importantly, we found that the reinstatement of contingency-discriminating patterns during memory retrieval was predictive of trial-by-trial performance; these patterns were stronger before correct, compared to incorrect, behavioural responses to tone presentation (Fig. 2f). This contingency-selective pattern reinstatement was not related to a firing rate bias of member neurons or animal running speed (Supplementary Fig. 2j,k). Conversely, the activation strength of contingency-invariant patterns was not related to memory performance (Fig. 2f). These findings suggest that, while contingency-invariant patterns are rapidly computed in the hippocampal CA1 at the onset of unsupervised exploration of a novel environment, spiking activity is further organised to form additional patterns which are sensitive to the learning contingencies experienced in that environment and predictive of behavioural performance.

During exploration of a novel environment, the coincidental spike discharge of CA1 neurons with spatially-overlapping firing fields gives rise to spatially tuned coactivity^5,12^. To test whether hippocampal CA1 follows such an inside-the-field (i.e. infield) firing association rule during contingency learning, we computed spatial maps both from the activation timecourses of the detected patterns, and from the spike trains of each of their member neurons (Fig. 3a and Supplementary Fig. 3). Contingency-discriminating coactivity was markedly less spatially coherent than that of contingency-invariant patterns (Fig. 3a,b). This was concomitant with less spatially coherent firing of contingency-discriminating pattern members relative to their contingency-invariant counterparts (Supplementary Fig. 4a,b). Moreover, individual firing member fields within the same contingency-discriminating pattern were less spatially congruent than those of a given contingency-invariant pattern (Fig. 3a,c; Supplementary Fig. 3). This was unrelated to differences in temporal correlation amongst members’ spike trains (Supplementary Fig. 4c). In fact, while contingency-invariant coactivity was spatially biased towards the place fields of member neurons, this bias was significantly weaker for contingency-discriminating patterns (Fig 3d,e). Thus, while congruent, contingency-invariant coactivity patterns represent space, contingency-discriminating coactivity is spatially untuned, consistent with a specialisation in representing ongoing behavioural contingency regardless of the animal’s location.

**Figure 3.**
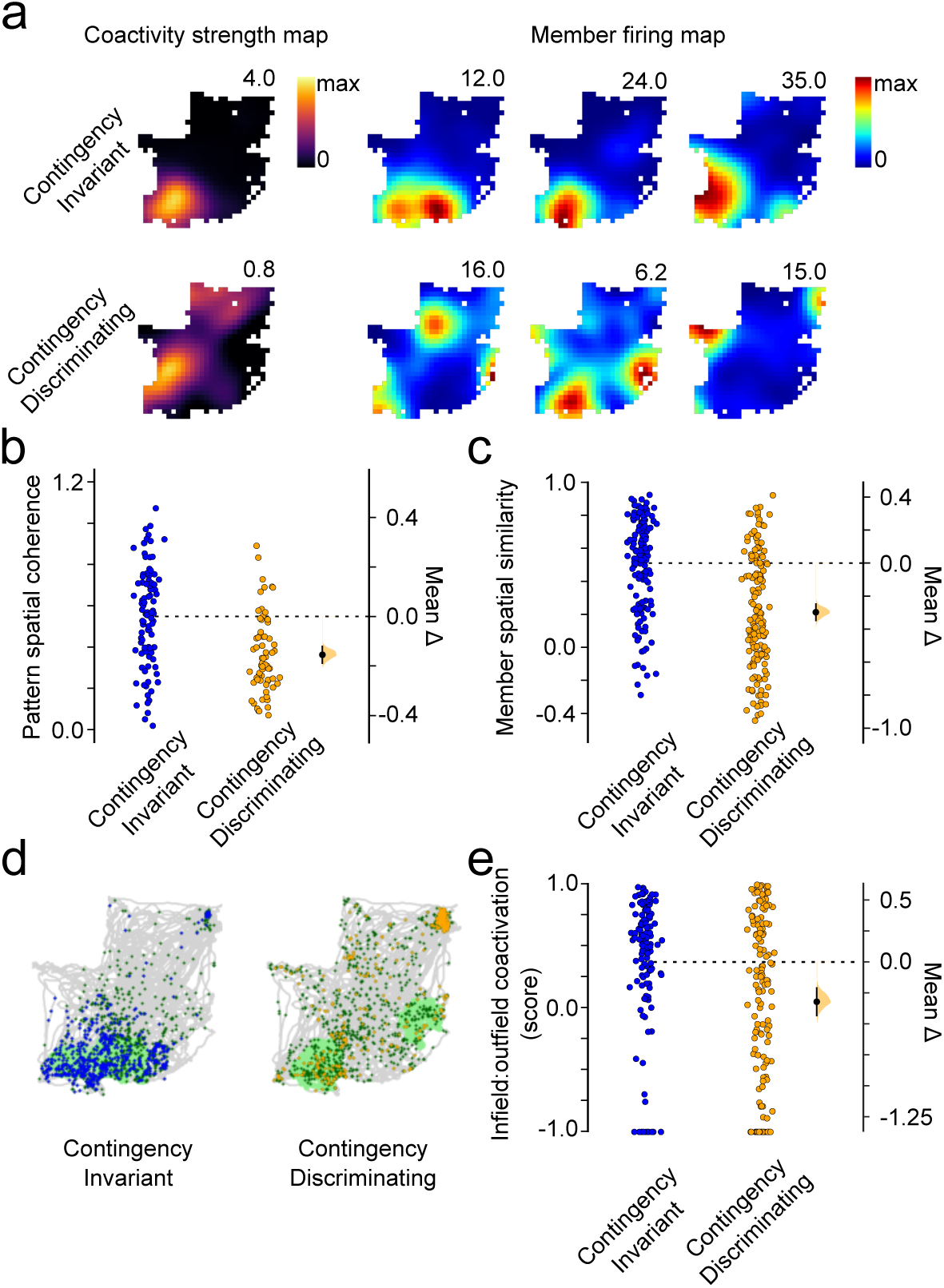
Contingency-discriminating CA1 patterns are spatially discontiguous and incongruent with their member neurons. **a)** Example coactivity pattern strength maps (left) and corresponding firing rate maps (right) of member neurons for a contingency-invariant (top row) and a contingency-discriminating (bottom row) pattern. **b)** Contingency-discriminating coactivity is less spatially coherent than that of contingency-invariant patterns. **c)** Member neuron firing fields are less spatially overlapping for contingency-discriminating than contingency-invariant patterns. **d)** Example coverage traces (grey) with overlaid spiking activity (dots) of a member of a contingency-invariant (left) and one of a contingency-discriminating (right) coactivity pattern. Spikes during a co-activation event of a given pattern are marked in blue (contingency-invariant) or orange (contingency-discriminating), while the remaining spikes are marked in green. Spatial firing field indicated by the green shading. **e)** Infield versus outfield co-activation score for member neurons of contingency-invariant and discriminating patterns.

Finally, we asked what circuit mechanisms support the organisation of CA1 spiking into task-related coactivity patterns during contingency learning. CA1 coactivity could rely on the recurrently connected upstream CA3 area^13,14^. Moreover, we previously found robust synaptic plasticity of left, but not right, CA3 (CA3^L^) inputs to CA1^15,16^. Converging evidence implicates such synaptic plasticity in mediating associative learning^17,18^. We hypothesised that plastic CA3^L^ inputs are instrumental in the gradual development of contingency-discriminating CA1 coactivity patterns during learning. To test this hypothesis, we transduced CA3^L^ pyramidal neurons of Grik4-cre mice with the yellow light-driven proton pump Archaerhodopsin-3.0 (Fig. 4a,b); implantation of tetrodes and optic fibres further allowed simultaneous monitoring of, and light delivery to, CA1 neuronal ensembles. Light delivery to CA3^L^ axons in CA1 during learning markedly reduced the power of theta-nested slow-gamma, but not mid-gamma, oscillations in CA1 (Fig. 4c, Supplementary Fig. 5a,b), consistent with the suggestion that CA1 slow-gamma oscillations report incoming CA3 inputs^19–21^. Yet, selectively suppressing CA3^L^→CA1 input preserved both the average firing rate of CA1 pyramidal neurons and the proportion of CA1 neurons assigned to coactivity patterns (Supplementary Fig. 5c,d). Importantly however, CA3^L^→CA1 input suppression shifted the distribution of CA1 between-contingency coactivity pattern similarity, and coactivity strength, towards contingency-invariance (Fig. 4d, Supplementary Fig. 5e). Moreover, CA3^L^→CA1 input suppression biased coincidental spike discharge towards member place fields (Fig. 4e), consistent with a loss of the spatially incongruent coactivity that characterises contingency-discriminating patterns. At the behavioural level, suppressing CA3^L^→CA1 inputs during learning had no effect on ongoing learning performance (Supplementary Fig. 5f) but impaired memory during the subsequent probe, selectively when mice had to flexibly retrieve two contingencies (Fig. 4f), but not when retrieving only one contingency (Supplementary Fig. 5g). Moreover, flexible memory retrieval was preserved after suppressing the more stable inputs from the right hemisphere (CA3^R^→CA1 inputs; Supplementary Fig. 5h-j). Together, these findings show that plastic CA3^L^ inputs support the organisation of CA1 spiking into contingency-discriminating coactivity patterns and associated two-contingency memory.

**Figure 4.**
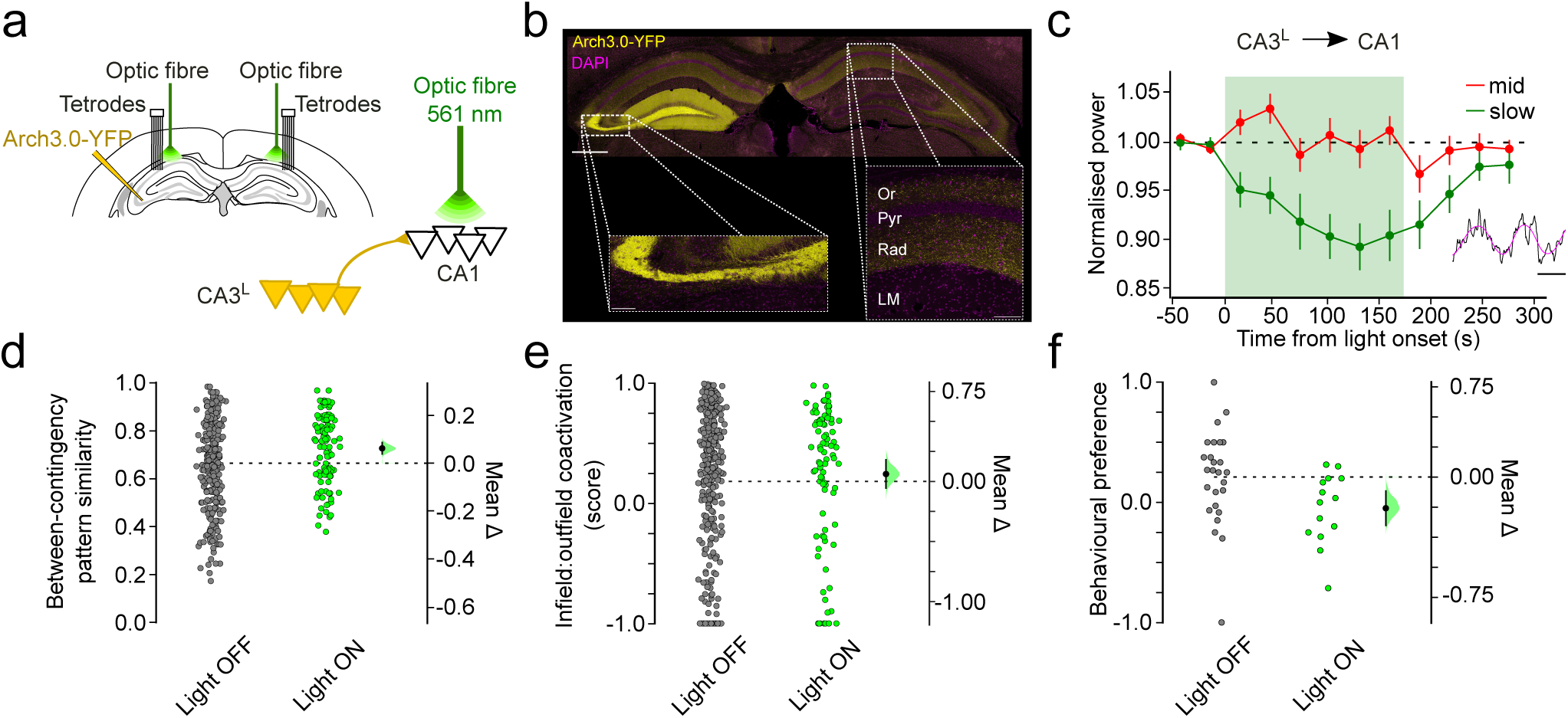
CA3^L^ inputs are necessary for contingency-discriminating CA1 coactivity and dynamic two-contingency memory. **a)** CA3^L^→CA1 optogenetic silencing protocol. CA3^L^ neurons were transduced with Archaerhodopsin 3.0 in Grik4-Cre mice (n=5) and their axonal projections in the CA1 targeted bilaterally during learning with yellow 561nm-light delivery from implanted optic fibres; 12 tetrodes monitored CA1 neurons. **b)** (Top) Expression of Archaerhodopsin3.0-YFP in the somata of CA3^L^ neurons and their axons in CA1 bilaterally; DAPI-stained nuclei. Higher-magnification images of YFP-expressing CA3 neurons (bottom left) and their axons in the contralateral CA1 (bottom right). Scale bars (top=100µm, bottom=10µm). Stratum: Or, Oriens; Pyr, Pyramidale; Rad, Radiatum; LM, Lacunosum Moleculare. **c)** Light delivery to Arch3.0-expressing CA3^L^ axons reduced the power of theta-nested slow, but not mid, gamma oscillations in CA1 (inset: example trace showing consecutive theta cycles nesting strong mid (∼50-90 Hz) and slow (∼25-40 Hz) gamma oscillations respectively. Raw trace and theta component as black and magenta traces, respectively; scale bar=100 ms). **d)** CA3^L^→CA1 input suppression shifted the between-contingency similarity of CA1 patterns towards contingency invariance. n=54 and 57 patterns detected in contingency *X* and contingency *Y* respectively on days with CA3^L^→CA1 input suppression (light ON days); 4.9±0.3 total neurons per pattern. **e)** Infield versus outfield co-activation score for member neurons of all patterns during CA3^L^→CA1 input suppression (light ON) versus light OFF days. **f)** CA3^L^→CA1 input suppression during learning impaired subsequent probe trial performance. Error bars, mean±S.E.M except when used with DABEST plots, where they represent mean±95% confidence intervals

Our findings identify a role for emergent, coactivity-based neural discrimination of behavioural contingencies in flexible memory-guided behaviour. The necessity of plastic inputs for discrimination at both the behavioural and neural levels reported here suggests an instrumental role for synaptic plasticity in the development of neural coactivity patterns that disambiguate new behavioural contingencies. This coding scheme could allow an efficient use of plasticity, generating a dynamic, coactivity-based discrimination of newly encountered contingencies every day, without committing individual neurons to represent such short-lived cognitive variables. A key prediction from this coding scheme is that downstream receiver neurons decode incoming information, represented as an emergent property of the collective activity of multiple neurons, by disambiguating relevant patterns of millisecond coincidence from the myriad of other inputs they receive^3^. In contrast, coactivity amongst congruent neurons would provide information redundancy for robust transmission to downstream neurons. Such redundancy could serve stable representation of the statistical regularities that define the background environment^22^. These contrasting coding schemes therefore provide distinct constraints on hippocampal communication with target regions involved in executive and motor functions. Altogether, our findings open new perspectives for future studies involving simultaneous ensemble recordings from the hippocampus and downstream target circuits to elucidate information decoding mechanisms.

## Figure Legends

**Supplementary Figure 1.**
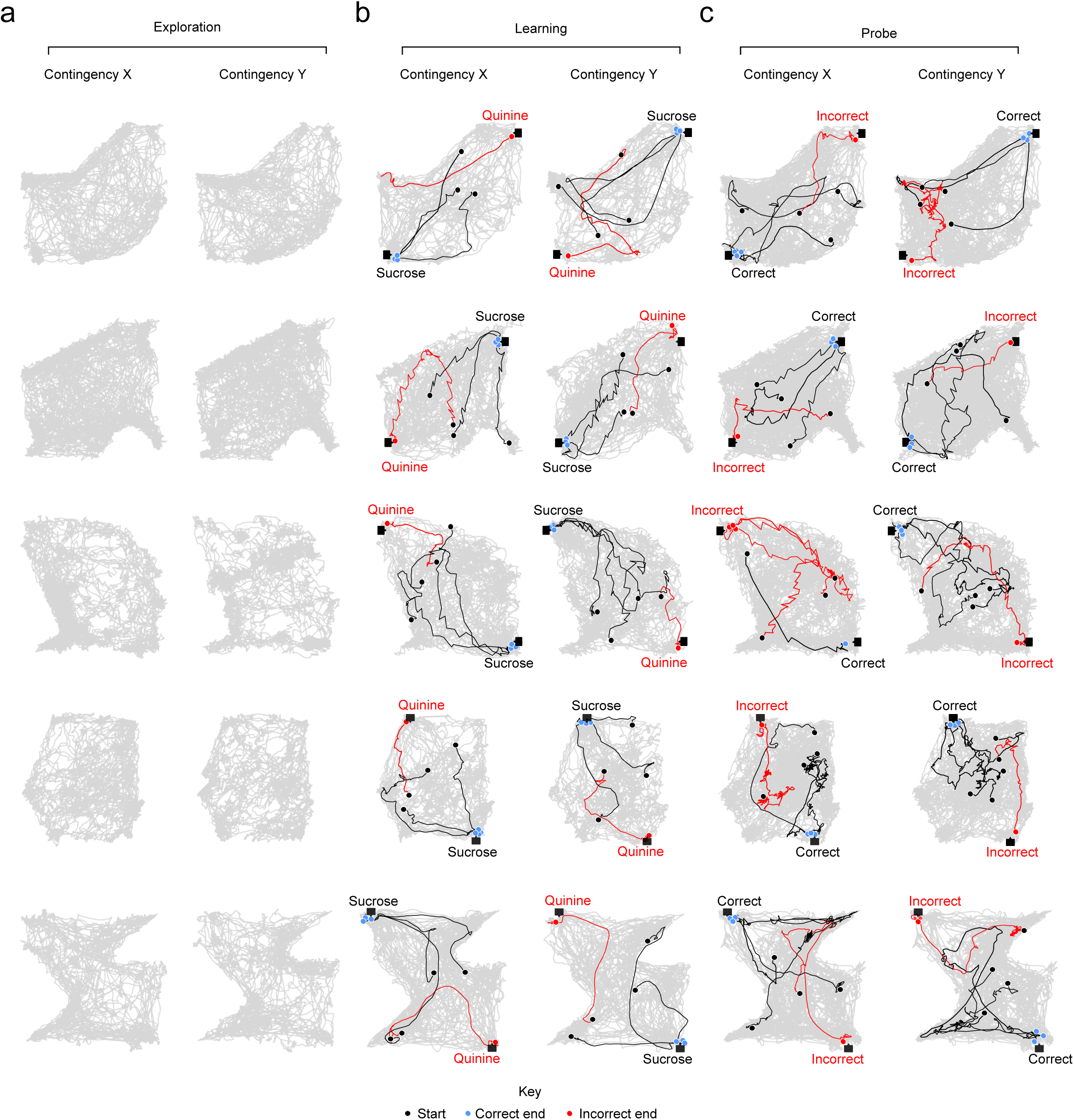
Enclosure set ups across distinct behavioural days. Animal coverage represented in grey. **a)** Example coverage paths for pre-learning exploration of learning enclosures. **b)** Example animal paths during learning trials in contingency *X* and contingency *Y*. **c)** Example animal paths during probe trials in contingency *X* and contingency *Y*. Paths of the animal during trials (correct path: black; incorrect path: red) are overlaid onto overall coverage (grey) for a single learning session. Black circles represent path starting points; blue and red circles represent correct and incorrect end points respectively.

**Supplementary Figure 2.**
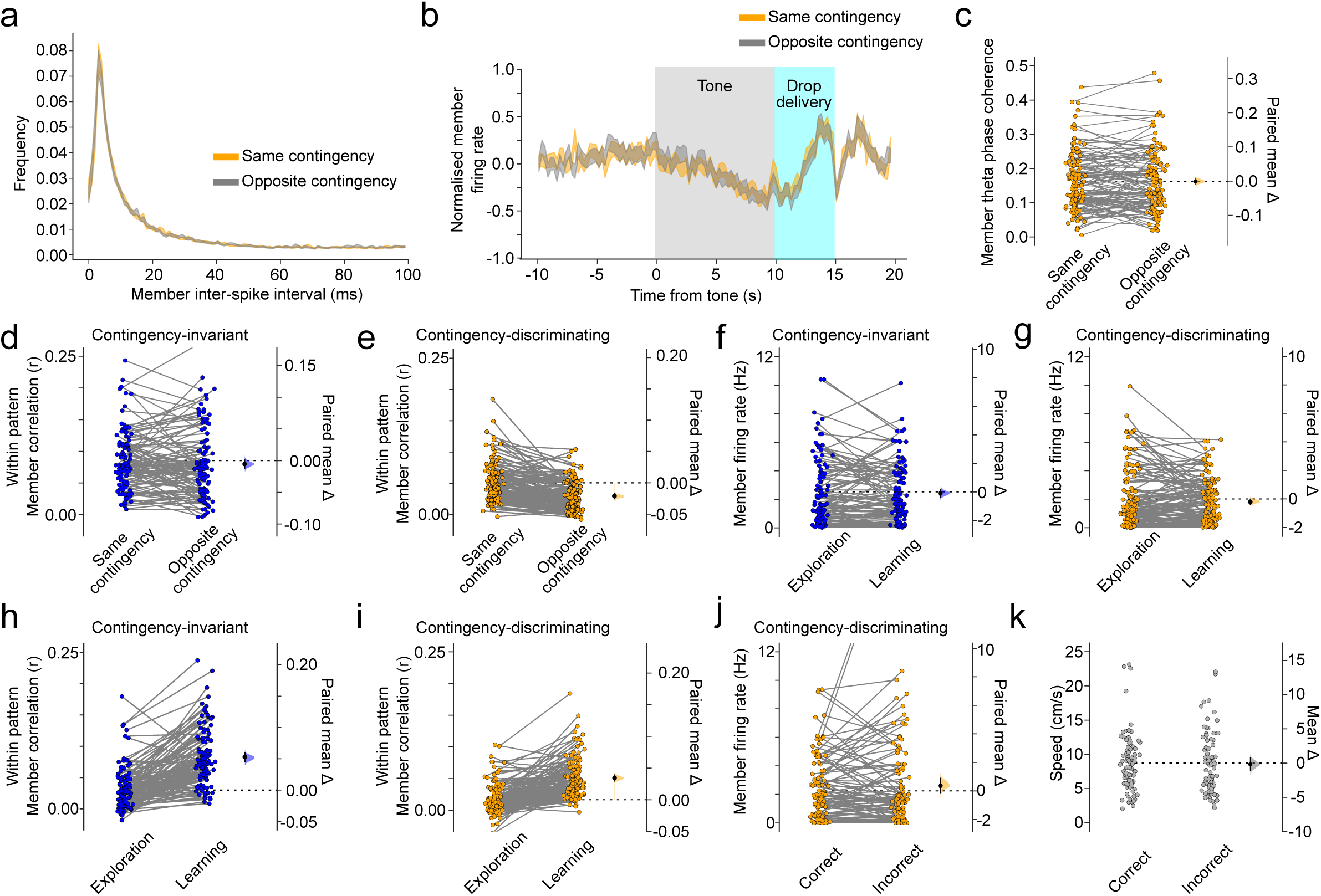
**a)** Inter-spike intervals, **b)** Z-scored firing rates during tone and drop delivery, and **c)** spike-phase coherence to theta oscillations of contingency-discriminating pattern member neurons in the same contingency (i.e. the contingency in which the patterns were detected) and the opposite contingency; colour coded orange and grey respectively. **d)** Mean temporal correlations amongst members of the same contingency-invariant and **e)** contingency-discriminating patterns in the same contingency and the opposite contingency. **f**,**g)** Mean firing rates and **h**,**i)** temporal correlations (Pearson r values) amongst each member neuron of a pattern and other members in the same pattern between learning and exploration. Firing rates of contingency-discriminating pattern member neurons remained indistinguishable between exploration and learning, while mean temporal correlations were higher in learning compared to exploration sessions. A similar result was seen for contingency-invariant pattern members (i.e. no change in rate but an increase in temporal correlation between exploration and learning). **j)** Contingency-discriminating pattern member firing rate is not higher before correct vs incorrect probe trials. **k)** Mouse running speed before correct and incorrect trials

**Supplementary Figure 3.**
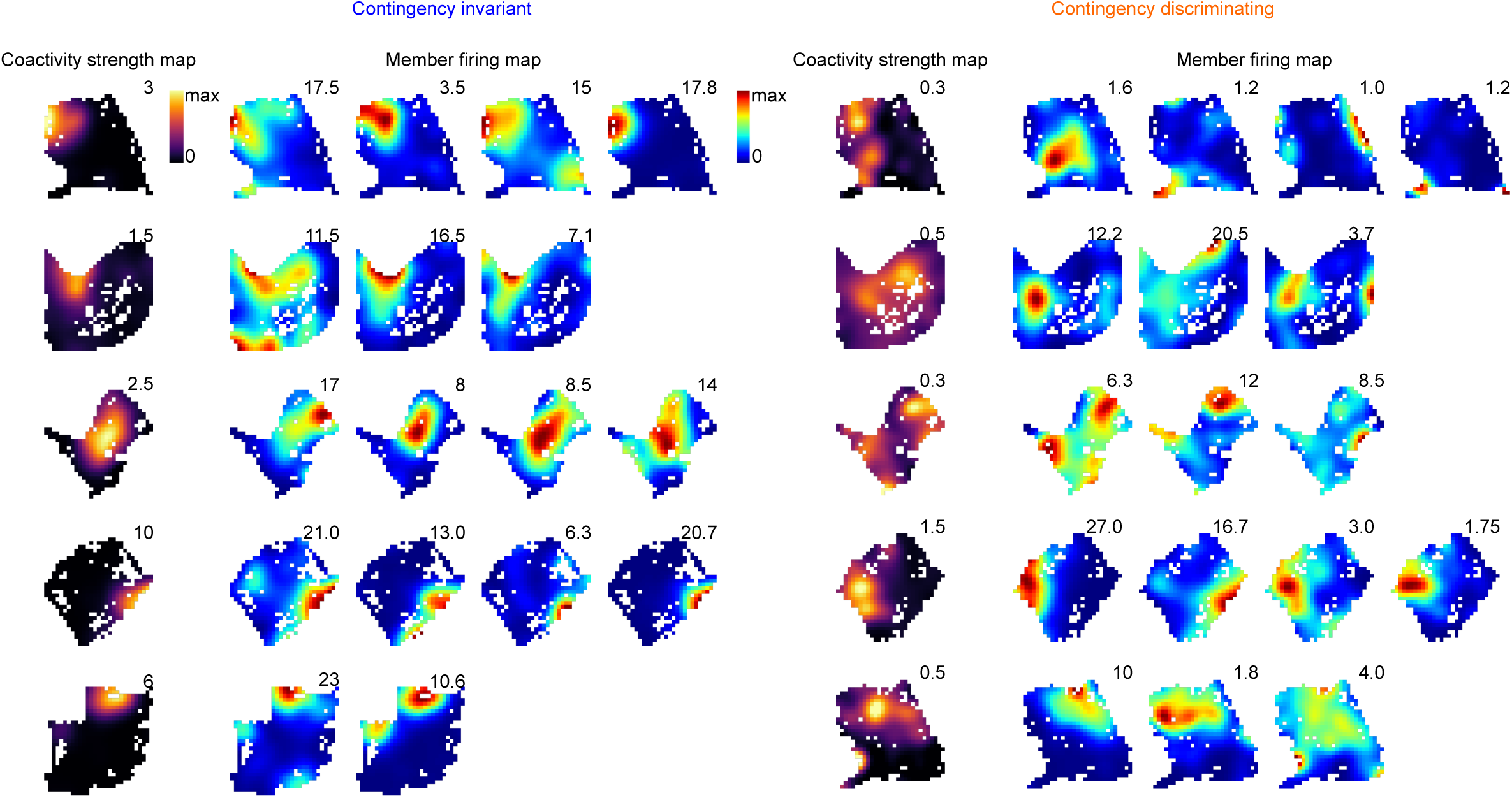
Example pattern activation maps and corresponding place maps of pattern member neurons for contingency-invariant (left) and contingency-discriminating (right) coactivity patterns.

**Supplementary Figure 4.**
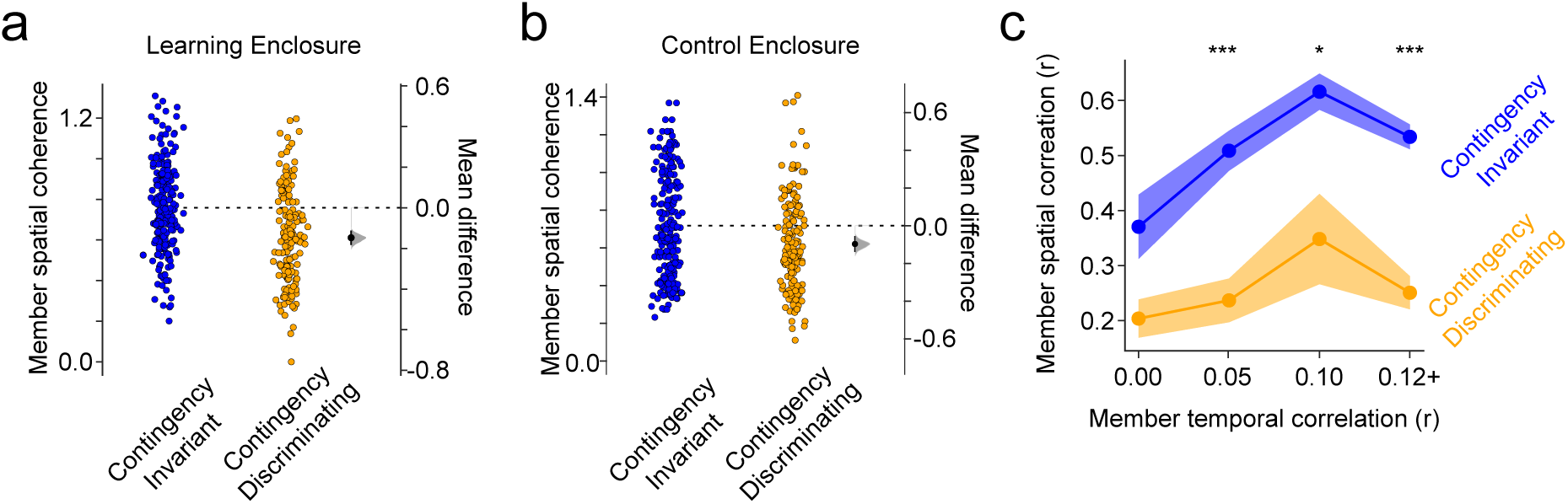
**a)** Spatial coherence of contingency-invariant is higher than that of contingency-discriminating pattern members in the learning and **b)** in the control enclosures. **c)** Spatial similarity of contingency-discriminating pattern members is lower than that of contingency-invariant pattern members regardless of the degree of temporal correlation between the member neurons. Two-way ANOVA with post-hoc Tukey-HSD test: main effect of pattern type (F=99.2, P<0.001) and temporal correlation (F=4.41, P=0.005). Pattern type: temporal correlation interaction (F=0.825, P=0.466).

**Supplementary Figure 5.**
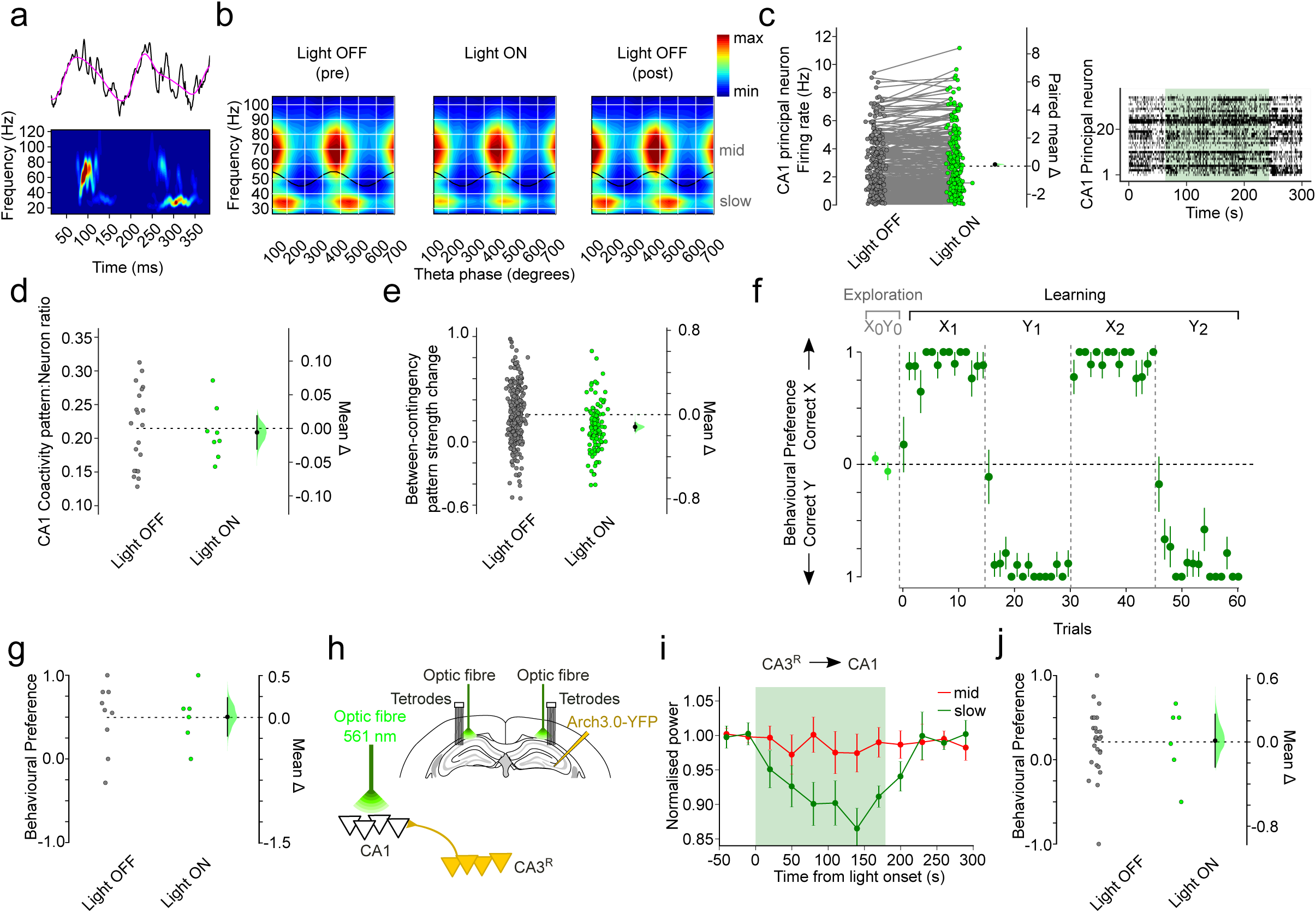
**a)** Example trace containing theta-nested mid and slow gamma oscillations (top; raw trace and theta component as black and magenta traces, respectively) along with its time-frequency representation (bottom) **b)** Example of the selective effect of CA3^L^→CA1 input suppression on the slow but not the mid gamma oscillations. Shown are Hilbert-spectra as a function of ongoing theta phase for pre, during and post light stimuli in a representative session. Theta cycles were subsampled to maintain instantaneous speed distributions across panels. **c)** Firing rate of CA1 principal neurons is unaltered by light delivery (*right*, example raster plot during light ON period for one recording day) **d)** The ratio of detected coactivity patterns to CA1 principal neurons is unaltered by CA3^L^→CA1 input suppression. **e)** The strength of coactivity patterns detected in the CA1 under CA3^L^→CA1 input suppression is markedly less sensitive to contingencies compared to light OFF days. **f)** CA3^L^→CA1 input suppression does not impair performance during learning trials. Mean correct dispenser preference; Light OFF: 0.87±0.10 (n= 42), Light ON: 0.87±0.10 (n=14); Unpaired two-sided t-test: P=0.98. **g)** Silencing CA3^L^ inputs to CA1 does not impair memory performance when each LED set signals the same contingency throughout all learning sessions (“Control days”). **h)** Schematic of CA3^R^→CA1 optogenetic silencing protocol. CA3^R^ neurons were transduced with Archaerhodopsin 3.0 in Grik4-Cre mice (n=3) and their axonal projections in the CA1 targeted bilaterally during learning trials with yellow 561nm-light delivery from implanted optic fibres. **i)** Suppression of CA3^R^→CA1 input also reduces the power of theta-nested slow gamma, but not mid gamma, oscillations. Three-way ANOVA with post-hoc Tukey-HSD test: Main effect of gamma type (F=9.59, P=0.003), light delivery (F=90.96, P <0.001), but no main effect of hemisphere (F=0.302, P=0.584) on normalised gamma power. Gamma type:light interaction (F=11.26, P=0.001). Post-hoc Tukey-HSD tests revealed that light had a significant effect on slow gamma power (P<0.001) but not mid gamma power (P=0.212), that this effect was significant for both hemispheres (P<0.001 for both left and right CA3→CA1 input suppression), and that normalised gamma power was indistinguishable with suppression of CA3 inputs from both hemispheres (P=0.721). **j)** CA3^R^→CA1 input suppression does not impair performance in probe trials. Error bars, mean ± s. e. m. except when used with DABEST plots, where they represent mean ± 95% confidence intervals. *** P<0.001, ** P<0.01, * P<0.05

## Methods

### Animals

We used male adult C57BL/6J mice (Charles River Laboratories; n = 3) and Grik4-cre mice^24^ (The Jackson Laboratory; n = 10). Animals were pre-selected based on their comprehensive coverage of a novel open field and approach to a sucrose-baited dispenser within this open field. Animals had free access to water in a dedicated housing facility with a 12/12 h light/dark cycle (lights on at 07:00h). Animals were housed with their littermates up until the start of the experiment. Food was available ad libitum before the experiments (see below), and water available ad libitum throughout. All experiments involving mice were conducted according to the UK Animals (Scientific Procedures) Act 1986 under personal and project licenses issued by the Home Office following ethical review.

### Surgical procedures

All surgical procedures were performed under deep anaesthesia using isoflurane (0.5-2%) and oxygen (2 l/min), with analgesia provided before (0.1 mg/kg vetergesic) and after (5 mg/kg metacam) surgery. For optogenetic manipulations, AAV5-EF1a-DIO-Arch3.0-YFP viral vector injections (2×500nl) were performed unilaterally in the dorsal CA3 on either the left or right hemispheres (CA3^L^ or CA3^R^) of hemizygous Grik4-cre mice using stereotaxic coordinates (site 1: −1.7 mm anteroposterior, ±1.5 mm lateral and −2.1 mm ventral from bregma; site 2: −2.3 mm anteroposterior, ±2.3 mm lateral and −2.3 mm ventral from bregma). The viral vector was delivered at a rate of 100 nl min^−1^ using a glass micropipette. For electrophysiological recordings, mice were subsequently implanted with a microdrive with 12-14 independently movable tetrodes (combined with two optic fibres for optogenetic manipulations; Doric Lenses) targeting the dorsal CA1 bilaterally^25^.

### Behaviour

After the recovery period of at least one week following the surgical implantation procedure, mice were familiarised daily to the experimental paradigm, including handling, connection to the recording system and exploration of open fields. Mice were maintained at 90-95% of their free-feeding bodyweight. Animals were made familiar with exploring an open field (“control enclosure”) and trained in three phases. Pre-training phase 1 involved conditioning mice to collect transiently available sucrose drops from a single dispenser following a ten second tone. Sucrose was initially available for 20 seconds before the drop was automatically aspirated by the dispenser. Over multiple training sessions, this was gradually reduced in 5 second intervals every time the mouse successfully obtained sucrose three times consecutively, until a 5 second availability period was reached. To encourage full coverage across the open field, and discourage persistence at the sucrose dispenser, tones were only delivered after the mouse had moved away from the dispenser to explore the open field. Training continued until mice successfully obtained reward on more than 80% of trials, while exploring the open field; this typically required 5-7 training days. All mice actively covered the open-field enclosures and approached the dispensers upon tone presentation. Next, for phase 2 of pre-training, mice experienced two pairs of wall-mounted LED displays and two identical dispensers in a novel spatial configuration of the learning enclosure each day. One dispenser delivered sucrose and the other quinine, with both drops simultaneously available for 5 seconds at the offset of a 10 second tone. The identity of the dispenser delivering each liquid was predicted by the currently illuminated, wall-mounted LED set, which defined a given contingency (*X* or *Y*). On a given phase 2 day, mice initially explored the control enclosure for one session, followed by the learning enclosure for two sessions, with a different LED display illuminated in each learning enclosure exploration session. Subsequently, a total of 6 learning sessions (3 of each contingency) were conducted in a pseudorandom order (e.g. *XYYXYX* or *XYXYYX*), with 15 tones (thus 15 trials) in each session. Sessions of the same contingency were never presented 3 times in a row. Sucrose and quinine were delivered simultaneously after 80% of the tones in each session, with the remaining 20% of tones being non-reinforced (no sucrose or quinine delivered). After at least 3 days (and up to 7 days) of training on phase 2, animals reached an average performance of 80% correct choices or more on a given phase 2 day and thus were ready for the training phase. All behavioural and electrophysiological data quantified here are from this training phase. Here, the procedure was almost identical to phase 2 except that only two sessions of each contingency were presented in alternation (XYXY) in a novel configuration of the enclosure each day; and a memory probe test was carried out one hour after the final learning session of the day. In the probe session, 24 trials were delivered under extinction (i.e. non-reinforced trials where neither sucrose nor quinine was available after the tone). 12 trials were presented in each LED-defined contingency, with pseudorandom transitions between *X* and *Y* LEDs while the animal was in the learning enclosure, with the restriction that either 2 or 5 trials would be delivered in succession while a given LED was on. On “Control days”, we tested learning and retrieval of a single behavioural contingency throughout all sessions. Here, task structure was identical to one-day learning except that the same dispensers delivered sucrose or quinine regardless of the currently illuminated LED display. In all cases, correct and incorrect trials were defined by which dispenser the animals approached during the 5 second period of reward delivery in learning trials, or during the period from the start of the tone up to 10 seconds after the tone during the probe. If more than one dispenser was approached during the probe, the first dispenser visited was used for scoring. Behavioural preference for correct (sucrose-delivering) and incorrect (quinine delivering) dispensers during learning and probe sessions (Fig. 1e,f) was calculated as the difference between correct and incorrect trials divided by the sum of the two trial types. A similar behavioural preference score was calculated for the exploration session, on the basis of visits to a dispenser instead of trials (since there were no trials during exploration; Fig. 1e).

### In vivo ensemble recordings and light delivery

On the morning of each recording day, optimal positioning within the CA1 pyramidal layer was carried out using the local field potential (LFP) signals obtained from each tetrode^25^ in search of multi-unit spiking activity. Tetrodes were then left in position for ∼1.5h before commencing recordings. Tetrodes were raised at the end of each recording day to avoid possible mechanical damage overnight. Optical interrogation was performed during learning using a diode-pumped solid-state laser (Laser 2000, Ringstead) that delivers yellow light (561nm; ∼5-7 mW output power) to the optic fibres implanted bilaterally above the CA1 pyramidal cell layer in order to suppress CA3→CA1 inputs in Arch3.0-expressing Grik4-cre mice. Mice were accustomed to light delivery before training. During training, light was delivered for 3 minute periods, 5 times per learning session, with a 2 minute light OFF gap between each light delivery. Trials occurred only during light ON epochs, and at least 1 minute after the light came on to allow sufficient time for axonal suppression^26^.

### Multichannel data acquisition

Amplification, multiplexing and digitisation of the signals from the electrodes was carried out using a single integrated circuit located on the head of the animal (RHD2164, Intan Technologies; gain x1000). The amplified and filtered (0.09Hz to 7.60kHz) electrophysiological signals were digitised at 20kHz and saved to disk along with the synchronisation signals from the position tracking and laser activation, as well as digital pulses reporting on tone presentation, sucrose and quinine delivery, drop removal, LED display illumination, and licking events at each dispenser (detected as a break in an infrared beam across each dispenser’s drop delivery port). To track the location of the animal, three LED clusters were attached to the electrode casing and captured at 25 frames per second by an overhead colour camera. The signal was transmitted offline and aligned with the registered analogue tracking position and the aforementioned digital pulse timestamps.

### Spike detection and unit isolation

The electrophysiological signal was subsequently band-pass filtered (800Hz to 5kHz) and single extracellular discharges were detected through thresholding the RMS power spectrum using a 0.2ms sliding window. Detected spikes of the individual electrodes were combined for each tetrode. To isolate spikes which putatively belong to the same neuron, spike waveforms were first up-sampled to 40kHz and aligned to their maximal trough. Principal component analysis was applied to these waveforms ±0.5ms from the trough to extract the first 3-4 principal components per channel, such that each individual spike was represented by 12 waveform parameters. An automatic clustering program (KlustaKwik, http://klusta-team.github.io) was run on this principal component space and the resulting clusters were manually recombined and further isolated based on cloud shape in the principal component space, cross-channel spike waveforms, auto-correlation and cross-correlation histograms. A fully-automated clustering was further performed using Kilosort (https://github.com/cortex-lab/KiloSort) via the SpikeForest sorting framework (https://github.com/flatironinstitute/spikeforest), with units then automatically curated using metrics derived from the waveforms and spike times, and verified by the operator. All sessions recorded on the same day were concatenated and clustered together. A cluster was only used for further analysis if it showed: stable cross-channel spike waveforms, a clear refractory period in its auto-correlation histogram, well-defined cluster boundaries and an absence of refractory period in its cross-correlation histograms with the other clusters. We isolated 997 principal neurons (731 in light OFF days and 266 in light ON days).

### Neuronal pattern isolation and tracking

Firing patterns of co-active CA1 principal cells were detected using an unsupervised statistical framework based on independent component analysis^11^. Spikes discharged by each neuron were counted in 25ms time bins and standardised (z-scored, i.e., the activity of each neuron was set to have null mean and unitary variance), to avoid *a priori* bias toward neurons with higher firing rates. The neuronal population activity was represented by a matrix in which each element represents the z-scored spike count of a given neuron within a given time bin. We extracted coactivity patterns from this matrix in a two-step procedure. First, the number of significant co-activation patterns embedded within the neuronal population was estimated as the number of principal components of the activity matrix with variances above a threshold derived from an analytical probability function for uncorrelated data. Second, we applied independent component analysis to extract the coactivity patterns from projection of the data into the subspace spanned by the significant principal components (i.e., each coactivity pattern was captured by an independent component). Pattern detection was performed using active periods (speed > 2 cm/s) separately during the entire last session of contingency X, contingency Y (i.e. X2 and Y2) or the exploration session of the control enclosure as appropriate. To assess the enclosure- or contingency-specificity of coactivity patterns, we compared all patterns detected across enclosures or contingencies, respectively. The similarity of two coactivity patterns was quantified using the absolute value of the cosine similarity of their normalised weight vectors. To quantify the degree of similarity between patterns detected in any two sessions, we identified the maximum similarity between a given coactivity pattern in one session and all patterns detected in the other. Contingency-discriminating patterns were defined as those for which maximum between-contingency similarity was below the 90^th^ percentile of the between-enclosure similarity distribution. A matched group of patterns at the top end of the between-contingency similarity distribution were used as “contingency-invariant” patterns for comparison. Subsequently, all patterns that had a within-contingency maximum similarity below the 90th percentile of the between-enclosure similarity distribution were excluded from further analysis. Since detected weight vectors were asymmetrical (Fig 2a) the direction where weights were highest was assigned positive weights, and principal CA1 neurons whose weight was positive and exceeded 2 standard deviations from the mean were identified as pattern ‘members’. In total, the analyses shown in Figures 2 and 3 included 67 contingency-discriminating patterns (159 member neurons), 51 contingency-invariant patterns (134 member neurons); all patterns detected in light OFF and light ON days were used in Figure 4.

The activation strength of each coactivity pattern at time t (Fig. 2a, Supplementary Fig. 2a) was computed as:

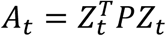

Where *Z*_*t*_ is a vector carrying the z-scored rate of each neuron at time *t, P* is the projection matrix (outer product) of the corresponding independent component, and *T* is the transpose operator. *A*_*t*_ is therefore the squared projection of *Z*_*t*_ onto the component that represents the coactivity pattern. This projection represents the similarity between the independent component and the population rate at a given time bin of 25 ms. The main diagonal of *P* was set to zero before calculating *A*_*t*_, in order to eliminate the contribution of single neurons to the coactivity pattern strength. The resulting value of *A*_*t*_ reflected expression strength of a particular coactivity pattern and was used in subsequent calculations of coactivity pattern emergence and spatial tuning. To determine whether pattern expression strength predicted probe performance, we calculated each pattern’s strength during the period of tone presentation but before the animal approached either dispenser, and averaged these values during theta cycles across epochs preceding correct or incorrect choices. The same calculation was performed for member neuron firing rates. Significant co-activation events (Figures 3 and 4) were defined as time points when co-activation strength was more than 2 standard deviations above the mean for the learning session in which the patterns were detected.

### Spatial maps

The recording arena was divided into bins of 1.5×1.5 cm to generate spike count maps (number of spikes fired in each bin) for each unit, or pattern strength map for each co-activation pattern, and an occupancy map (time spent by the animal in each bin). Rates and occupancy were calculated only during active periods (i.e. speed > 2 cm/s) and bins visited less than a total of 5 times per session were excluded from subsequent analysis. All maps were then smoothed by convolution with a two-dimensional Gaussian kernel of s.d. equal to two bin widths. Finally, spatial rate maps were generated by normalising the smoothed spike count maps by the smoothed occupancy map. Spatial coherence reflects the similarity of the firing rate in adjacent bins, and is the z-transform of the Pearson correlation (across all bins) between the rate in a bin and the smoothed rate of the same bin^27^. The same calculation was used on coactivity pattern strength to calculate pattern spatial coherence. Spatial similarity between maps of member neurons, or co-activation patterns, was calculated as the Pearson correlation coefficient from the direct comparison of the spatial bins between the smoothed place rate maps. To determine the degree to which pattern coactivations were biased relative to members’ firing fields, we calculated an infield coactivation score for each member as the spatial density of coactivations inside the member neuron’s firing field (spatial bins within 70% of the peak firing rate bin) minus the outside-the-field coactivation density divided by the sum of those two values.

### LFP analyses

Raw local field potentials (LFPs) were down-sampled from 20kHz to 1250 Hz (order 8 Chebyshev type I filter was applied prior to decimation to avoid aliasing) and then decomposed using Empirical Mode Decomposition (EMD^28^). In order to avoid mode mixing, we used the mask sift EMD procedure^29^, with sinusoidal masks with the following frequencies: 350, 200, 70, 40, 30 and 7 Hz, which captured mid, slow gamma and theta oscillations as isolated components. To determine individual theta cycles and theta phase, we first detected peaks and troughs of theta with absolute values higher than low-frequency component (sum of all components with main frequencies below the theta signal) envelope; then a theta cycle was defined by pairs of supra-threshold troughs separated at least by 71ms (∼14 Hz) and no more than 200ms (5 Hz) that surrounded a supra-threshold peak. Theta phase was calculated by linear interpolating neighbouring theta troughs, zero crossings and peaks. For nested-gamma analysis (Fig. 4c and Supplementary Fig. 5a,b,i), instantaneous envelopes and frequencies were calculated by means of the normalised-Hilbert transform^28,30^. For the time course analysis shown in Fig. 4c and Supplementary Fig. 5i, we adopted a bootstrap procedure to keep their speed distribution of each time bin virtually equal. For each experiment, we used 60 to 30-second pre-laser stimulus windows as a reference for speed distribution. More specifically, we calculated the histogram (linearly-spaced speed bins from 2 to 30 cm/s) of instantaneous speed values for each theta cycle within that reference window; then a bootstrap consisted of (1) subsampling theta cycles from that reference time window by randomly choosing 75% of the cycles in each speed bin (i.e., maintaining the original speed histogram proportions); (2) from all remaining time windows, for each theta cycle in the reference window we randomly chose a cycle with matched speed (no more than 2.5% away from the reference cycle). One hundred of such bootstraps were computed for each tetrode, then all tetrodes of each experiment were averaged. Figures show the mean across recording days.

### Anatomical and histological analysis

All mice were anaesthetised with pentobarbital following completion of the experiments and transcardially perfused with PBS followed by 4% PFA / 0.1% glutaraldehyde in PBS solution. Brains were extracted and kept in 4% PFA for at least 24 h before slicing. Coronal sections (50 μm thick) were then made and stored in PBS-azide combined with DAPI to stain neuronal somata. All sections were mounted in Vectashield (Vector Laboratories, Cat. No. H-1000) and images of native eYFP fluorescence and DAPI fluorescence were captured with a LSM 710 (Zeiss) confocal microscope using ZEN software (Zeiss).

### Statistical analyses

Error bars, mean±S.E.M except when used with DABEST plots, where they represent mean ± 95% confidence intervals. Ns refer to recording days for behavioural preference figures and theta-nested gamma analysis. For multi-unit data, Ns refer to coactivity patterns, coactivity pattern members or all principal neurons as indicated. All *P* values were calculated using a two-sided t-test, or ANOVA followed by a *post hoc* Tukey’s HSD test for multiple comparisons; unless specified otherwise. Data Analysis with Bootstrap-coupled Estimation (DABEST^23^ plots are used throughout the manuscript to plot data against a mean (or paired mean) difference between the two conditions (right y-axis) and compare this difference against zero using the bootstrapping generated 95% confidence intervals (error-bar).

## Acknowledgements

We thank Alexander Morley for developing the automated spike sorting software used here. We also thank Stephen McHugh, Stephanie Trouche and David Bannerman for helpful advice when designing the behavioral protocol; Jane Westcott and Ben Micklem for technical support; Andrew J. Quinn for developing the EMD toolbox we used here; Helen Barron and all the members of the Dupret laboratory for inspiring discussions. This work was supported by the Biotechnology and Biological Sciences Research Council UK (Award BB/N0059TX/1) and the Medical Research Council UK (Programme MC_UU_12024/3).

